# Role of periodic forcing on the stochastic dynamics of a biomolecular clock

**DOI:** 10.1101/2020.12.03.410530

**Authors:** Zhanhao Zhang, Supravat Dey, Abhyudai Singh

## Abstract

Biomolecular clocks produce sustained oscillations in mRNA/protein copy numbers that are subject to inherent copy-number fluctuations with important implications for proper cellular timekeeping. These random fluctuations embedded within periodic variations in copy numbers make the quantification of noise particularly challenging in stochastic gene oscillatory systems, unlike other non-oscillatory circuits. Motivated by diurnal cycles driving circadian clocks, we investigate the noise properties in the well-known Goodwin oscillator in the presence and absence of a periodic driving signal. We use two approaches to compute the noise as a function of time: (i) solving the moment dynamics derived from the linear noise approximation (LNA) assuming fluctuations are small relative to the mean and (ii) analyzing trajectories obtained from exact stochastic simulations of the Goodwin oscillator. Our results demonstrate that the LNA can predict the noise behavior quite accurately when the system shows damped oscillations or in the presence of external periodic forcing. However, the LNA could be misleading in the case of sustained oscillations without an external signal due to the propagation of large noise. Finally, we study the effect of random bursting of gene products on the clock stochastic dynamics. Our analysis reveals that the burst of mRNAs enhances the noise in the copy number regardless of the presence of external forcing, although the extent of fluctuations becomes less due to the forcing.

## I. INTRODUCTION

Biomolecular clocks are gene regulatory networks in cells that can produce sustained oscillations in gene products to keep precise time for the various process such as cell cycle dynamics, maintenance of daily-cycle in most organisms, and vertebra formation in mammals [1]–[7]. Behind each biomolecular clock, there is a nonlinear gene regulatory network with varying complexity that generate sustained oscillations [4], [7]–[9]. A common mechanism for sustained oscillations in various clock such as circadian clocks, segmentation clocks, and cell cycle dynamics is time-delayed auto-inhibition [7]–[9]. In the case of the circadian clock, besides the autonomous oscillations, there is periodic signaling due to daily light-dark cycle that couples and synchronizes the clock with the environment [5], [6]. In the case of segmentation clocks, adjacent cells are coupled via signaling molecules to maintain synchrony with neighbors for perfect vertebra formations [7], [10], [11].

The expression of a gene is subject to fluctuations due to intrinsic and extrinsic factors [12]–[19]. Inherent stochasticity of biochemical reactions is the source of intrinsic noise. Whereas environmental fluctuations and other cell-specific differences such as cell-cycle stages, cell sizes, availability of enzymes for mRNA and protein synthesis are the major sources of extrinsic noise [20]–[23]. Depending on the contexts, the noise can have consequences that could be beneficial or detrimental to the cells [24]–[28]. The fluctuations are even present in the case of oscillatory gene expressions [10], [29], [30]. How biomolecular clocks keep precise time amid such fluctuations is an intriguing fundamental question.

Several experimental [10], [31]–[34] and theoretical [7], [35]–[38] studies in the past have studied the role of intercellular and external coupling on the synchronization the rythms in the gene expression in various genetic circuits. However, in the presence of randomness, the role of coupling on the synchronization is not well understood. In this paper, we investigate the effect of coupling to the external environment on the stochastic dynamics of a biomolecular clock. In the presence of a periodic signal, we study the well known Goodwin oscillator based on time-delayed autoinhibition regulatory circuit where the signaling effect is incorporated in the synthesis rate of the clock gene mRNA. We find that the periodic forcing can entrain the clock dynamics, and the entrainment patterns depend on the frequency of amplitude of the signal. The exact analytical approach for solving noise in the oscillatory gene expression is challenging as the circuits consist of nonlinear elements. Here, we first write down the first- and second-order moment dynamics using the linear noise approximation (LNA) that linearizes nonlinear propensities, assuming small fluctuations around the mean. Then numerically solve them to compute the noise. By comparing the LNA results with the exact stochastic simulations, we find the prediction of LNA results is quite well when the system shows damped oscillations or in the presence of external periodic forcing. However, the LNA results could be misleading for sustained autonomous oscillations (without external signal) where large fluctuations also propagate in time. Finally, we study the effect of bursty production of mRNA. We observe that mRNA bursts enhances the noises in the copy numbers, regardless of the presence of external forcing. Although, the extent of fluctuations become less due to the forcing.

### Notation

We use regular lower case letters to denote deterministic variables and bold lowercase letters for stochastic variables. For example, *m*(*t*) represents the level of mRNA at time *t* in deterministic analysis, and ***m***(*t*) represents mRNA count in stochastic case. For stochastic case, the angular bracket 〈·〉 is used to represent the ensemble averages of a quantity. For example, 〈***m***(*t*)〉 and 〈***m***(*t*)^2^〉 are the mean and second moment of the mRNA count at time *t*.

## II. Deterministic dynamics of the Goodwin oscillator in the presence of periodic forcing

To investigate the role of external periodic forcing on the deterministic and stochastic dynamics of an autonomous biomolecular clock, we formulate a model using the well known Goodwin oscillator [39]. Below, we briefly discuss the deterministic dynamics of the Goodwin oscillator before proceeding to the incorporation of the periodic external signal.

### A. Dynamics without periodic forcing

The deterministic dynamics of the Goodwin oscillator, involving three species namely an mRNA *M*, intermediate protein *E*, and a repressor protein *P*, is given by the following ordinary differential equations (ODEs) [39], [40],

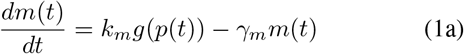

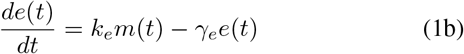

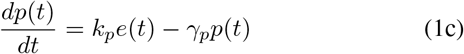

Here, *m*(*t*), *e*(*t*), and *p*(*t*) denote the concentration of the species *M, E*, and *P* at time *t*. The molecular mechanism behind the dynamics is as follows (see Fig. 1): the gene produces mRNA *M*, mRNA is then translated into intermediate protein *E, E* activates the production of the repressor protein *P*, and finally, *P* represses the mRNA synthesis and closes the negative feedback loop. The parameter *k_m_* is the maximum synthesis rate for *M, k_e_* and *k_p_* are the synthesis rates of *E* and *P*, and *γ_m_, γ_e_*, and *γ_p_* are the degradation rates. The repression is the Hill-type and given by *g*(*p*) as

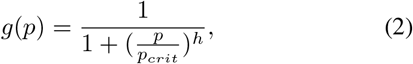

where the parameter *h* is the Hill coefficient and *p_crit_* is the value of the concentration required for the half maximal repression.

Here we note that the dynamics of intermediate species creates a time delay for the repression for the mRNA production and essential for the generation of sustained oscillations. Besides, the repression must be sufficiently strong with the Hill coefficient *h* > 8 [40]. For *h* < 8, the system can show damped oscillations, not sustained one.

**Fig. 1.**
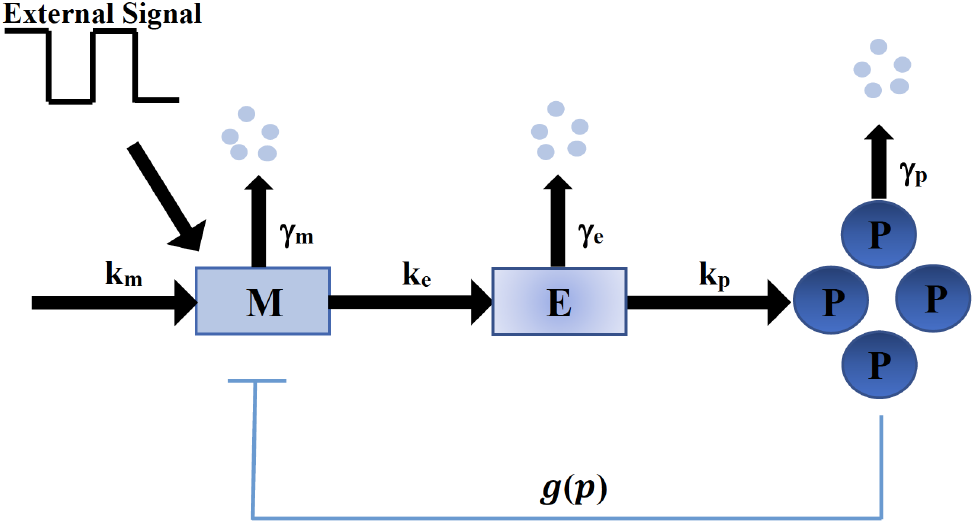
A schematic of the model. The regulatory network for the Goodwin oscillator in the presence of external periodic signal: The network consisting of three species that create delayed negative feedback for mRNA synthesis. The mRNA *M* is produced from a gene and then translated into the intermediate protein *E*. The protein *E* then actives the production of the repressor protein *P*, which inhibits the productions of *M* and closes the negative loop. The dynamics of *E* makes a delay for the repression. Finally, the autonomous Goodwin oscillator is coupled to a periodic external signal that modifies the mRNA synthesis rate.

### B. Incorporation of periodic forcing

Here, we consider the Goodwin oscillator in the external periodic signal. The external signal could be dark-light cycle or temperature cycle as in the case of circadian clocks. We assume that the external periodic signal only effects the synthesis rate of clock gene mRNA linearly. Therefore, the modified mRNA dynamics in the presence of external signal Φ(*t*) is given by,

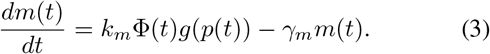

The dynamics for the concentration of *E* and *P* proteins is assumed to be unaltered by the signal and follow (1b and 1c). We consider a square waveform for the external signal with amplitude *A_s_* and frequency *f_s_*. We numerically solve the above system for a given set of parameters and present our results below.

### C. Entrainment Behavior

The frequency of the autonomous oscillators *f*_0_ depends on the parameter values. The typical sustained oscillations in the mRNA level in the absence forcing is shown in Fig. 2 (A). In the presence of periodic forcing, the mRNA level and its frequency adjust themselves to follow the signal as shown in Fig. 2 (B)-(D). This phenomena is known as entrainment. We observe the entrainment behavior for wide ranges of amplitude *A_s_* and frequency *f_s_* variation. Different entrainment patterns, known as ‘Arnold’s tongue’, emerge as *A_s_* and *f_s_* are varied. For example, for a small *A_s_* the system shows 1:1 entrainment (one clock oscillation peak for one signal peak) when the *f_s_* is close to *f*_0_. However, for a large *A_s_*, 1:2 is observed (one clock oscillation peak for two signal peaks). This type entraiment behavior is very common in gene oscillations circuits [32]–[35].

**Fig. 2.**
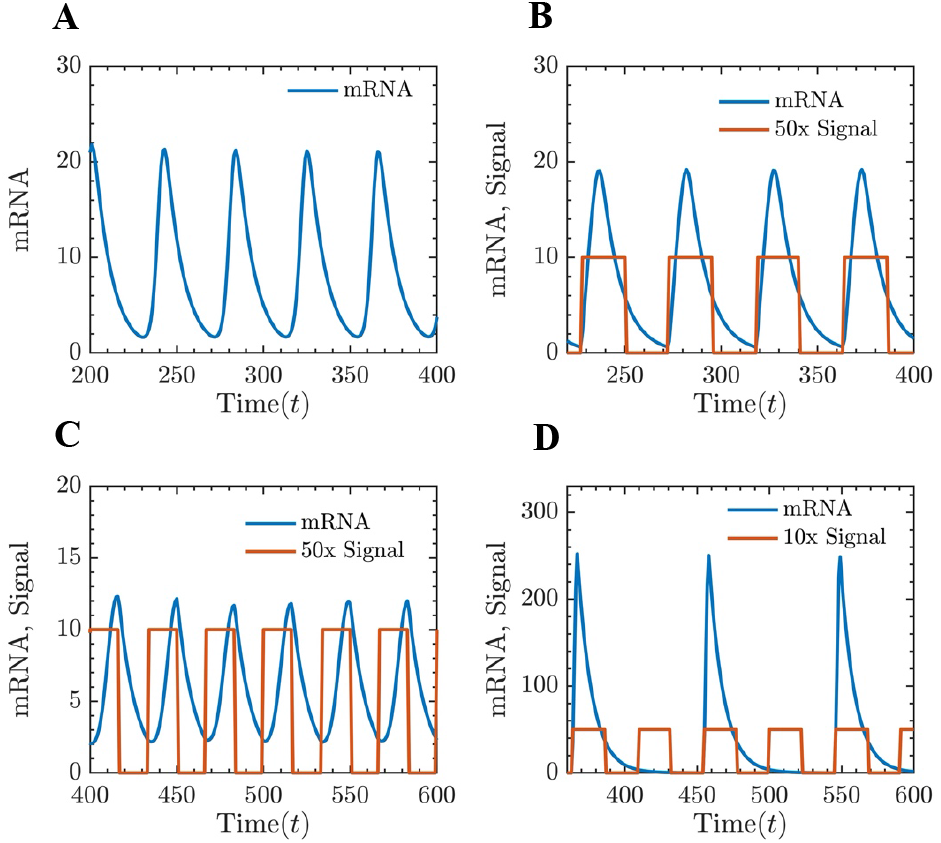
Entrainment due to periodic forcing. Typical trajectories for mRNA concentration of autonomus Goodwin oscillator in the presence and absence of forcing. (**A**) The sustained oscillations in the absence of external forcing (with *f*_0_ ≈ 0.025). (**B** and **C**) For a small signal amplitude (*A_s_* = 0.2), a 1:1 entrainment is observed when the signal frequency is close to the autonomous as in B for *f_s_* ≈ 0.9 *f*_0_ and in C for *f_s_* ≈ 1.2 *f*_0_. (**D**) A different entrainment pattern can be observed when *A_s_* is large even though *f*_0_ and *f_s_* are close. For amplitude *A_s_* = 1 and *f_s_* ≈ 0.9 *f*_0_, a 2:1 entrainment is observed. Parameter for Goodwin oscillator: *k_m_* = 20, *k_e_* = 1, *k_p_* = 0.2, *γ_m_* = *γ_e_* = *γ_p_* = 0.1, *p_crit_* = 100, and *h* = 12.

## III. Stochastic Dynamics

In the above section, we have studied entrainment for the deterministic case. However, the copy numbers of mRNAs and proteins inside genetically identical cells, even under the same external environment show large fluctuations (gene expression noise). These fluctuations are inevitable as they arise due to biochemical reactions that are inherently stochastic and occur in low molecular copy numbers, regardless of nonoscillatory [12]–[19] or oscillatory gene expressions [10], [29], [30]. Here, we investigate entrainment behavior in the presence of inherent stochasticity.

In our stochastic setting, all the biochemical reactions for the productions and degradations occur probabilistically. We also want to understand how the external signal controls the noise that arises from the bursty production of mRNA. Bursty productions are sources of high gene expression noises. Gene expression bursts have been experimentally observed in diverse cell types [41]–[46], and correspond to distinct mechanisms at the transcriptional and translation level. The production of mRNA as a bursty process, which can be thought of as a result of stochastic switching between multiple promoter states [43], [47]–[49]. In this analysis, we assume that the copy numbers of mRNA released in a burst follow the (shifted) geometric distribution

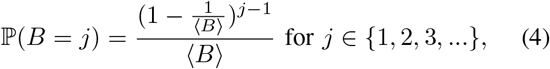

where 〈*B*〉 is the average burst size. The other productions are assumed to follow simple birth process.

Let us denote the stochastic variables for the copy numbers of the mRNA, intermediate protein, and repressor protein at time *t* by ***m***(*t*), ***e***(*t*), and ***p***(*t*). Then, the transition probability of all the synthesis and degradation reactions in the time interval (*t, t* + *dt*] are given by,

Bursty mRNA synthesis:

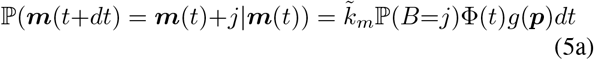

mRNA degradation:

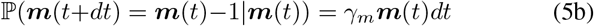

Intermediate protein synthesis:

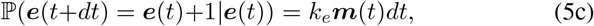

Intermediate protein degradation:

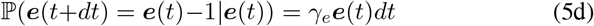

Repressor protein synthesis:

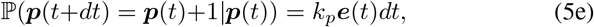

Repressor protein degradation:

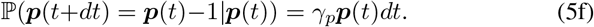

Here, 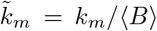 is the scaled burst frequency which cancels the burst effect on the mean dynamics. The set of transition probabilities define the dynamics of stochastic system of the Goodwin oscillator in the presence of external signal. To get the dynamics in the absence of periodic signal, Φ(*t*) in (6a) must be replaced with 1. Let *P*(***m, e, p, t***) be the probability density for observing ***m*** copies of mRNA, ***e*** copies of intermediate protein, and ***p*** copies of the repressor protein at time *t*. Then the time evolution of *P*(***m, e, p, t***) is given by the chemical master equation (CME) [50]

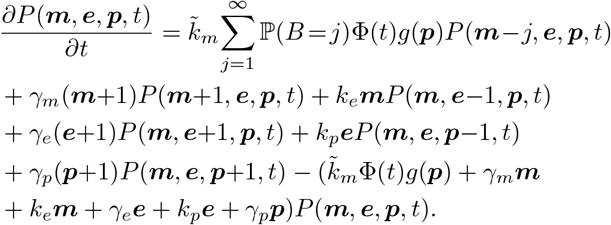

### A. Moment dynamics and the LNA

The nonlinearity in the above equation makes it impossible for solving it for the full probability distribution. Rather, we are interested in computing the noise in copy numbers as a function of time which relies on the first two moment dynamics. Therefore, we want to derive the dynamical equation for the first two moments. The dynamics of any arbitrary moment 〈***m**^n1^ **e**^n2^ **p**^n3^*) where *n1, n2, n3* ∈ {0,1,2,..} for the above stochastic systems can be obtained from the CME and is given by the Dynkin’s equation [51],

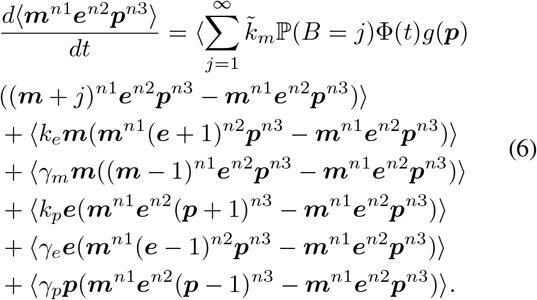

The nonlinearlity in the propensity of the mRNA synthesis due to the function *g*(***p***) makes the moment dynamics unclosed where the dynamics of a lower order moment depends on the higher moments [51]. We use the Linear Noise Approximation(LNA) to obtain close moment dynamics [50], [52]–[54]. Under the LNA, we linearize the *g*(***p***) assuming the fluctuation around the mean 〈(***p***)〉 is small:

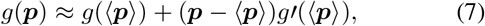

where *g*′(***p***) is the derivative of *g* with respect to ***p***. Using (7), we obtain the first and second order moment dynamic equations of the system from the above Dynkin’s equation. Here we list them:

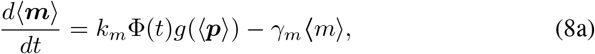

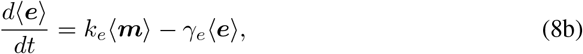

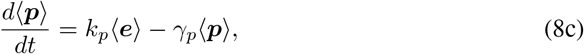

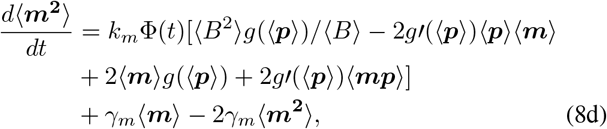

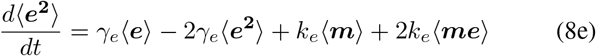

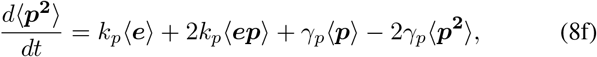

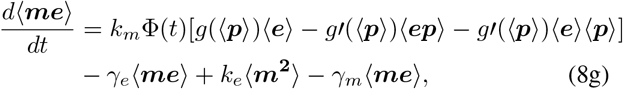

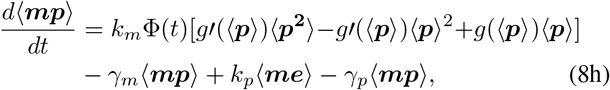

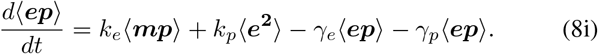

Please note that under this approximation, the first order moments dynamics (9a–c) reduces to the deterministic system discussed before. By numerically solving these 9 ODEs (8), we obtain values of 〈***m***^2^〉 and 〈***m***〉 to compute the noise level, considering zero species at *t* = 0. We quantify the noise in mRNA count as function of time in terms of the Fano factor *F_m_*(*t*):

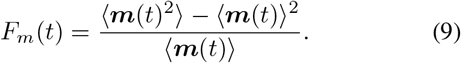

### B. Exact Stochastic Simulations using Gillespie algorithm

We perform stochastic simulations to check the validity of the LNA. We evolve the stochastic system (5) in numerically exact way using Gillespie algorithm [55] using our own code written in C++. According to this algorithm, at a given time *t*, we choose a reaction randomly according to its transition probability from (5). Then we update the species counts accordingly and increment the time by a random amount that is drawn from an exponential with whose mean is the total transition rate (sum of transition rates for all the events) of the system at that time. We generate a large number of stochastic trajectories starting from zero species at *t* = 0 and compute the noise statistics.

### C. Results

#### Damped oscillator in the presence of periodic forcing

We first study how an external periodic signal entrains a damped oscillation dynamics. We note that for a Hill coefficient *h* < 8, the Goodwin system show damped oscillations. Fig. 3(A) shows a typical trajectory for a damped system for *h*=4 in the absence of periodic forcing. The mean (〈***m***〉) and the noise (*F_m_*(*t*)) dynamics for the mRNA counts obtained from both the stochastic simulations and LNA are shown in Fig. 3(C), (E), respectively. Both the mean and noise show oscillating modulations that eventually dies out and converges to fixed values for large *t*. The match between the stochastic simulation and the LNA is very good.

**Fig. 3.**
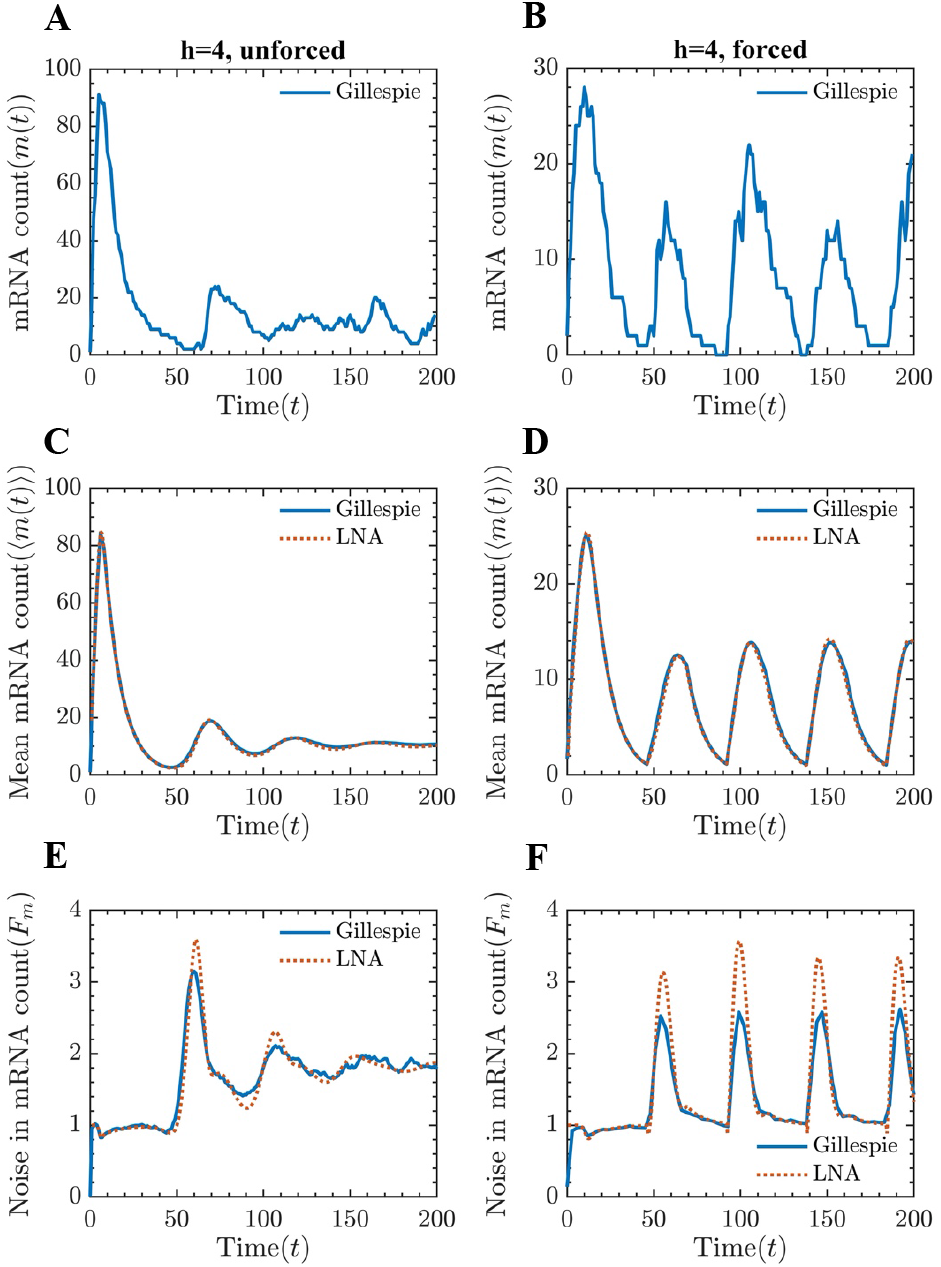
A damped system in the absence and presence of periodic force: (**A**) and (**B**) Typical stochastic trajectories for the mRNA count for the damped oscillator in the absence and presence of external signalling. (**C**) and (**D**) The mean level of mRNA computed using the LNA and Gillespie are plotted against time in the presence and absence of external signalling. The external force can drive damped oscillations to sustained oscillations. The match between the LNA is excellent. (**E**) and (**F**) The Fano factor obtained from the LNA and stochastic simulations are plotted as a function of time. The quantitative match between LNA and Gillespie is quite good, except for noise values near the peaks in the presence of forcing. Parameters: *k_m_* = 20, *k_e_* = 1, *k_p_* = 0.2, *γ_m_* = *γ_e_* = *γ_p_* = 0.1, *p_crit_* = 100, *h* = 4, *A_s_* = 0.2, *f_s_* = 0.022*Hz* and 〈*B*〉 = 1.

In the damped system, periodic forcing can generate sustained oscillations that get entrained with the signal (see Fig. 3(B)). The LNA and stochastic simulation results for the mean and noise match quite well (see Fig. 3(D), (F)). The noise *F_m_*(*t*) also shows sustained oscillations. The peak in the noise and the mean level do not appear at the same time point, but they do appear in the proximity. The quantitative match can be poor near the peaks of the oscillating noise.

#### Sustained oscillator in the presence of periodic forcing

The trajectories of the sustained oscillations in the absence and presence of external signal are shown in Fig. 4(A) and (B), respectively. As can be seen, the external signal makes the clock oscillations more precise. Interestingly, the mean behavior of stochastic simulations and LNA results show distinct behavior (see Fig. 4(C) and (D)). While the LNA (the same as deterministic) results shows sustained oscillations, the Gillespie results seem to follow a damped dynamics in the absence of external signal (Fig. 4(C)). This damped behavior at the mean level obtained from Gillespie simulations is as a result of the loss of coherence due to large stochasticity in the absence of entrainment. In the presence of external forcing, however, both LNA and Gillespie results show sustained oscillations (see Fig. 4(C)), as entrainment makes the clock more precise in time. The quantitative match between LNA and Gillespie results are very good, except near the peaks(see Fig. 4(D)).

**Fig. 4.**
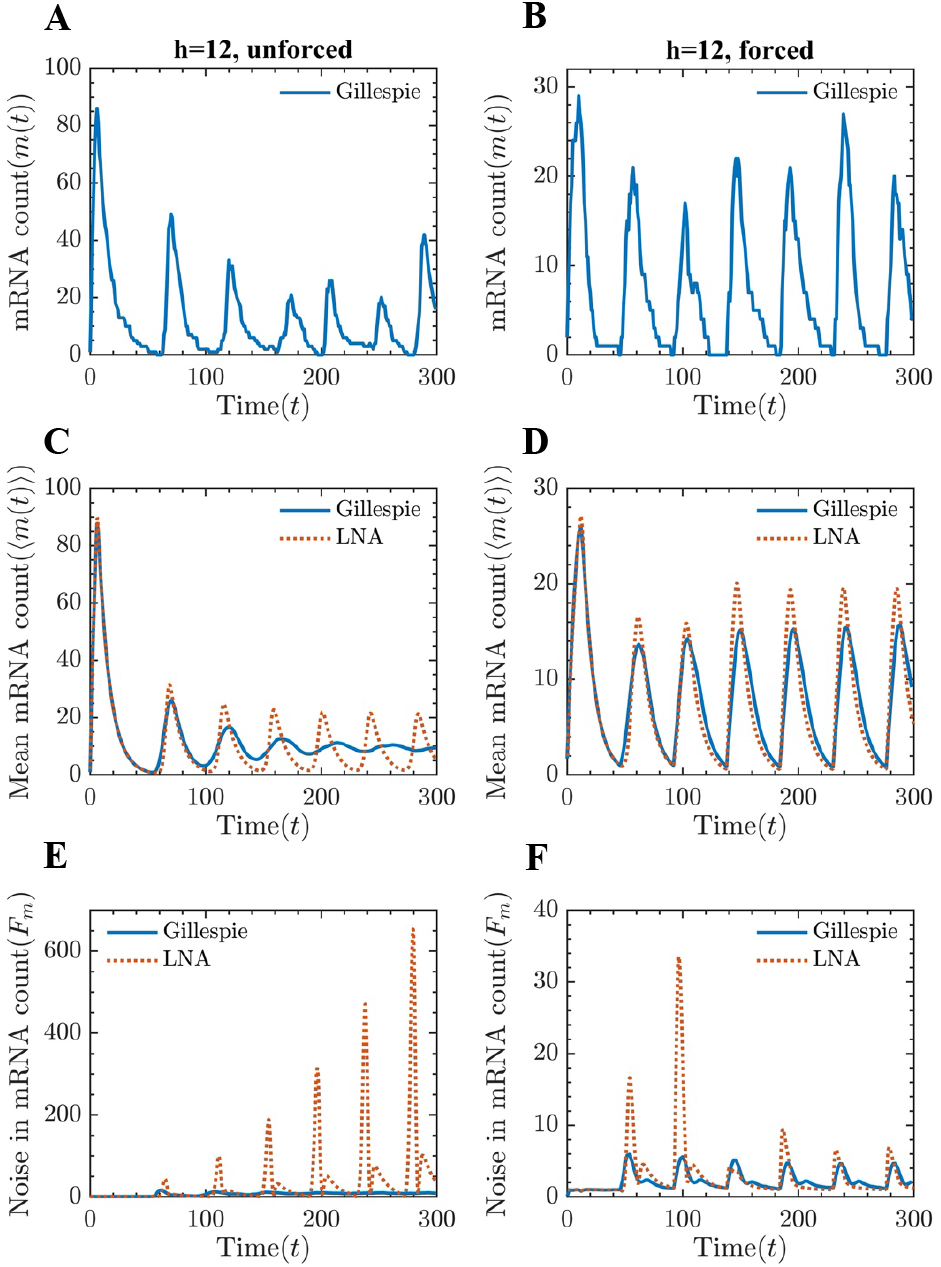
An autonomous system in the absence and presence of periodic force: (**A**) and (**B**) Typical stochastic trajectories showing sustained oscillations for the mRNA count in the absence and presence of external signalling. (**C**) and (**D**) The mean level of mRNA computed using the LNA and Gillespie are plotted against time in the presence and absence of external signalling. In the absence of external forcing, the mean trajectory for mRNA obtained using the stochastic simulations, show damped behavior inconsistent with the LNA results that predict sustained oscillations. In the presence of forcing, the match between the LNA and stochastic simulations quite good. (**E**) and (**F**) The Fano factor obtained from the LNA and stochastic simulations are plotted as a function of time. While the Gillespie results predict noise convergence, the LNA predicts divergent behavior as a function of time in the absence of periodic forcing. In the presence of periodic forcing, the noise from both the methods show sustained periodic modulations with good quantitative match for large time, expect peak regions. Parameters: *k_m_* = 20, *k_e_* = 1, *k_p_* = 0.2, 〈*B*〉 = 1, *γ_m_* = *γ_e_* = *γ_p_* = 0.1, *p_crit_* = 100, *h* = 12, *A_s_* = 0.2, and *f_s_* ≈ 0.9*f_0_*.

The noise behavior from the LNA is completely misleading in the absence of the periodic signal (Fig. 4(E)). The noise shows oscillating behavior that diverges with time. Again, this behavior arises because of the presence large fluctuations that propagate in time where the LNA breaks down. For the entrained case, the noise also shows sustained oscillations (Fig. 4(F)). The quantitative match is poor near the peaks at the early times.

To understand the effect of periodic forcing on the reg-ularity of the oscillations, we compute the distribution of the time period from the steady-state trajectories. In Fig. 5(A), we plot the distributions in the presence and absence of the periodic forcing. As can be seen, forcing makes the distribution narrower, causing more precise oscillations. We also compute the steady-state distributions in the peak values (Fig. 5(B)), and as expected, the distribution become narrower in the present of forcing.

**Fig. 5.**
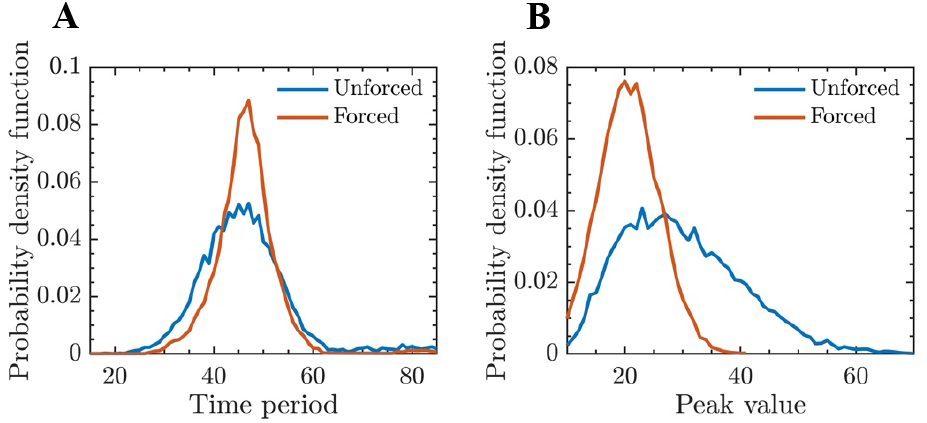
Stochastic simulation results for probability distri-butions of the time period and peak value. (**A**) The time period distributions in the presence and absence of periodic forcing. (**B**) The peak value distributions in the presence and absence of periodic forcing. The periodic forcing makes the distributions for time period and peak value narrower. Parameters: *k_m_* = 20, *k_e_* = 1, *k_p_* = 0.2, 〈*B*〉 = 1, *γ_m_*=*γ_e_*=*γ_p_*=0.1, *p_crit_*=100, *h* = 12, *A_s_* = 0.2 and *f_s_* ≈ 0.9*f*_0_.

### D. Effect of the burst sizes

In the no feedback limit, one it can be shown that the noise for bursty productions is given by its average burst size [56]. How does burst size affect the oscillatory circuit in the presence of periodic forcing? To address this issue, we study the effect of bursty production of mRNA with various burst sizes. All the results presented above were for with average mRNA burst size 〈*B*〉 = 1. Here, we consider the first peak value of the noise as a representative measure. However, the conclusion is independent of the specific choice. We plot the first peak noise value as a function of noise for damped and sustained systems in the absence and presence of periodic forcing in Fig. 6. The noise linearly increases with burst sizes for all the case. In the case of damped oscillation, the forcing does not alter the noise (Fig. 6(A)). However, in the sustained case, the forcing substantially reduces the noise.

**Fig. 6.**
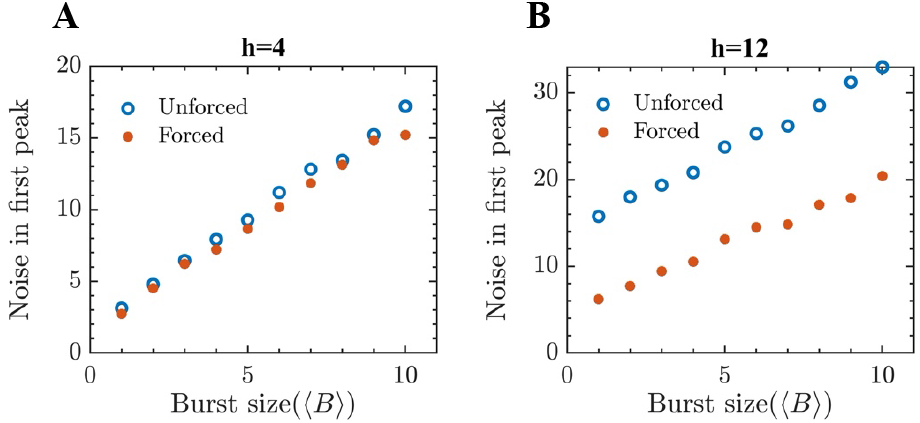
The relationship between burst sizes and the first peak value of Fano factor: (**A**) The noise increases with the burst size linearly and the noise in the presence of forcing is the same as no forced case. (**B**) Noise in unforced system in much larger than forced for sustained system, but both noises increase linearly with the burst size. Parameters: *k_m_*=20, *k_e_* = 1, *k_p_*=0.2, 〈*B*〉 = 1, *γ_m_*=*γ_e_*=*γ_p_*=0.1, *p_crit_*=100, *A_s_* = 0.2 and *f_s_* ≈ 0.9*f*_0_.

## IV. Conclusions

In summary, in this paper, we have investigated the effect of periodic forcing on the stochastic dynamics on a biomolecular clock, motivated by the circadian clock in the presence of the light-dark cycle. For this, we have considered the classic Goodwin oscillator in the presence of a squarewave signal. The presence of the external signal makes the clock precise in terms of amplitude and time period fluctuations. We have quantified the noise in the mRNA level as a function of time using two approaches: (i) solving moment dynamics obtained using the linear noise approximation, assuming small fluctuations around the mean, and (ii) using exact stochastic simulations. We have demonstrated that the LNA works well in the presence of periodic forcing when the fluctuations become smaller. For an autonomous oscillator, the LNA results can be misleading in the absence of periodic forcing due to large fluctuations in the time period and amplitude. Finally, we have studied the effect the mRNA bursts and have found entrainment is unable to reduce the effect of bursts.

## ACKNOWLEDGMENT

This work is supported by the National Science Foundation Grant ECCS-1711548.

## Notes

### Competing Interest Statement

The authors have declared no competing interest.

